# Applying joint graph embedding to study Alzheimer’s neurodegeneration patterns in volumetric data

**DOI:** 10.1101/2023.01.11.523671

**Authors:** Rosemary He, Daniel Tward, the Alzheimer’s Disease Neuroimaging Initiative

**Affiliations:** Departments of Computer Science and Computational Medicine, University of California, Los Angeles; Departments of Computational Medicine and Neurology, University of California, Los Angeles

**Keywords:** Alzheimer’s disease, structural MRI, graph embedding, network analysis, familywise error rate control

## Abstract

Neurodegeneration measured through volumetry in MRI is recognized as a potential Alzheimer’s Disease (AD) biomarker, but its utility is limited by lack of specificity. Quantifying spatial patterns of neurodegeneration on a whole brain scale rather than locally may help address this. In this work, we turn to network based analyses and extend a graph embedding algorithm to study morphometric connectivity from volume-change correlations measured with structural MRI on the timescale of years. We model our data with the multiple random eigengraphs framework, as well as modify and implement a multigraph embedding algorithm proposed earlier to estimate a low dimensional embedding of the networks. Our version of the algorithm guarantees meaningful finite-sample results and estimates maximum likelihood edge probabilities from population-specific network modes and subject-specific loadings. Furthermore, we propose and implement a novel statistical testing procedure to analyze group differences after accounting for confounders and locate significant structures during AD neurodegeneration. Family-wise error rate is controlled at 5% using permutation testing on the maximum statistic. We show that results from our analysis reveal networks dominated by known structures associated to AD neurodegeneration, indicating the framework has promise for studying AD. Furthermore, we find network-structure tuples that are not found with traditional methods in the field.

## 1 Introduction

Alzheimer’s disease (AD) is a progressive mental disorder associated with neurodegeneration that generally occurs in old ages. It is one of the most common diseases in seniors, killing more than breast cancer and prostate cancer combined(Association, 2019). However, AD can only be formally diagnosed through an autopsy after a patient is deceased, encouraging the research of alternative proxies for AD diagnosis. There are currently three classes of potential biomarkers that could offer useful alternatives to assess AD diagnosis: *β*—amyloid(A), tau(T) and biomarkers for neurodegeneration or neuronal injury(N)(Jack et al., 2016), with the last class being the focus of this paper. Neurodegeneration can be measured noninvasively, such as through structural MRI, but is not specific to AD as it can reflect other diseases(Jack Jr and Holtzman, 2013). For example, atrophy is a biomarker that is often associated with AD, but also occurs in a variety of disorders such as epilepsy and anoxia(Jack Jr and Holtzman, 2013). The lack of specificity of these neurodegeneration biomarkers poses a major limitation in their utility for early-stage clinical diagnosis of AD. A promising approach to improving specificity is to consider patterns of volumetric changes across the whole brain, rather than focusing on a small number of regions. For example, in discussing the diagnosis of Mild Cognitive Imparment due to AD, Albert et al. suggests the possibility of biomarkers describing “complex patterns of tissue loss” through “data driven statistical approaches in which many different brain regions are evaluated simultaneously” (Albert et al., 2011).

While rare in volumetric analysis, such patterns have been studied extensively in functional brain imaging. One popular approach to studying brain connectomics is to extract information about interactions between volumetric pixels (voxels) from time-series functional magnetic resonance imaging (fMRI) data collected over a timescale of weeks, months or years(Cohen et al., 2017). Functional connectivity, in particular, studies temporal dependencies among anatomically separated regions(Van Den Heuvel and Pol, 2010). There are several approaches to study functional connectivity in fMRI data. As our interests lie within the whole brain instead of a single voxel, we discuss only multivoxel pattern analysis methods that study networks as a whole(Lewis-Peacock and Norman, 2014). A traditional method in the field is seed-based analysis, in which a region of interest (ROI) is selected, and all voxels correlated to the ROI is identified(Cole et al., 2010). For example, in an earlier work Biswal et al. studied the motor cortex to identify the sensorimotor network(Biswal et al., 1995). Unsupervised clustering methods including k-means, hierarchical and graph-based methods do not require a priori ROI and group voxels together by their similarities in time series data(Khosla et al., 2019). In their paper, Lee et al. uses fuzzy-c-means clustering to identify resting state networks(Lee et al., 2012).

Over the years, graph-based approaches have gained popularity in studying functional connectivity. To convert a brain into a graph, the regions are modeled as nodes and connections between regions as edges. Under this model, one can construct a matrix of all pairs of connections in the brain, known as the functional connectome(Fornito et al., 2016), of which decomposition or embedding methods can be applied to uncover latent variables. Independent component analysis decomposes data into linearly independent components, grouping brain regions into networks based on their voxel activation correlations(Calhoun et al., 2001). Non-negative matrix factorization is a dimensionality reduction method that forces non-negativity constraints on the components(Khosla et al., 2019). Some popular embedding methods include Adjacency Spectral Embedding (ASE)(Sussman et al., 2012), which embeds a single symmetric adjacency matrix using eigenvectors corresponding to the largest eigenvalues, and Laplacian Eigenmap (LE)(Belkin and Niyogi, 2003), which embeds a single graph-Laplacian matrix using its eigenvectors corresponding to the smallest nonzero eigenvalues. However, several limitations lie within these graph-based approaches. First, they embed one graph at a time, and combining individual embeddings across multiple graphs is not a straigh-forward task. Second, the results are difficult to interpret, and further analysis is required(Yang et al., 2020).

One technique for embedding multiple graphs at once is omnibus embedding, in which the matrices of multiple graphs are combined into one and embedding is done on the big matrix(Levin et al., 2017). However, the combined matrix is usually very large and require lots of computational power. Dictionary learning is another framework for uncovering low-dimensional embeddings across multiple graphs, allowing for group comparison. Drawing upon clustering and linear decomposition methods, this method allows for additional constraints to achieve better formed solutions(Abraham et al., 2013). In the work of D’Souza et al.(D’Souza et al., 2019), they use a dictionary learning method to model interactions between resting state functional MRI and behavioral data in Autism Spectrum Disorder. Their method finds shared dictionary elements across multiple graphs and a subject specific loading onto the elements, which are then used as inputs to a neural network for disease prediction(D’Souza et al., 2019). In the work of Wang et al.(Wang et al., 2019), they propose a joint graph embedding method to estimate a low dimensional embedding across multiple graphs and each graph’s projection onto that embedding, which we will discuss more in detail in section 2.2.

In this work, we shift our attention away from functional connectivity and propose a graphbased approach to study neurodegeneration using volumetric data. To our knowledge, this is the first time joint graph embeddings have been used to study volumetric brain data. Similar to methods above, we model the brain as a graph where a node represents a structure of interest and an edge represents a correlation in atrophy patterns between two structures(Xu, 2021). We then use a multigraph embedding technique to try and understand these patterns and uncover potential biomarkers. By applying multigraph embedding to study neurodegeneration, which spans a much longer timescale than fMRI measures, we hope to increase specificity of our results. In addition, we hope to increase the specificity of neurodegeneration biomarkers by modeling the dataset with a more complex pattern than existing approaches. Rather than looking at volume changes in each structure individually in the traditional mass univariate method(Pengas et al., 2012), we add complexity by characterizing pair-wise relationships between structures in the context of uncovered networks.

## 2 Material and methods

### 2.1 Data preprocessing

We obtain our data from Alzheimer’s Disease Neuroimaging Initiative (ADNI) database (adni.loni.usc.edu). The ADNI was launched in 2003 as a public-private partnership, led by Principal Investigator Michael W. Weiner,MD. The primary goal of ADNI has been to test whether serial magnetic resonance imaging (MRI), positron emission tomography (PET), other biological markers, and clinical and neuropsychological assessment can be combined to measure the progression of mild cognitive impairment (MCI) and early Alzheimer’s disease (AD). Specifically, we took the ADNI1 3Y1.5T Longitudinal FreeSurfer dataset(Wyman et al., 2013) prepared by University of California, San Francisco, comprised of 699 individuals in total. We selected 108 regions of interest common in studying neurodegeneration, excluding non-brain and whole-brain structures as we are interested in structures on a smaller scale. We selected a cohort in which each individual has at least 3 visits during the span of 3 years. For subjects with missing volumes, we forward filled in time. We model each individual’s brain as a graph, where each anatomical structure is a node and an edge exists between two nodes if they are highly correlated during neurodegeneration. For each individual in the cohort, we converted volumetric data into a correlation matrix of size 108 by 108 and then an adjacency matrix based on a threshold of 0.8 by absolute value, as shown in Fig. 1. For defining disease groups, the Clinical Dementia Rating (CDR) is referenced, which consists of 5 levels: 0 (None), 0.5 (Questionable), 1 (Mild), 2 (Moderate), and 3 (Severe)(Morris, 1991). We decided a threshold of <= 1 to separate the group into none/mild and severe cognitive impairment. After filtering and pre-processing, the cohort contains 494 individuals, specifically the none/mild group with 322 and the severe group with 172.

**Fig. 1.**
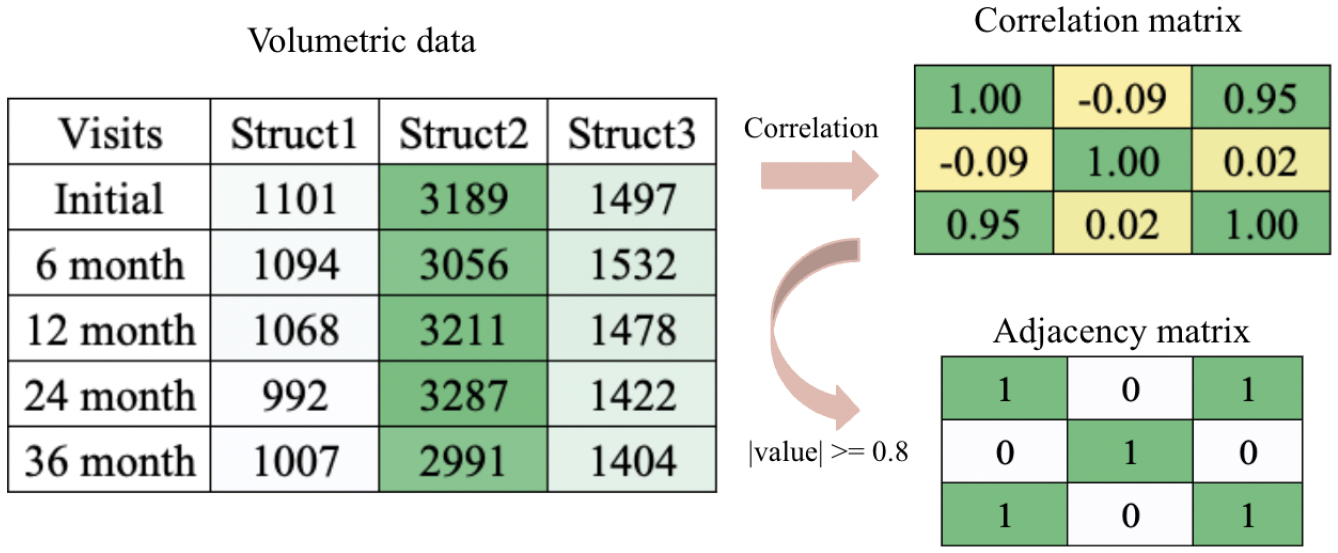
An example of transformation for one individual’s selected structures

### 2.2 Multiple Random Eigengraphs and Joint Graph Embedding

We first review the mathematical model and original joint graph embedding algorithm proposed by Wang et al.(Wang et al., 2019), of which our algorithm is based on. In this work, we refer to a random graph as a graph in which the edges are generated under a probability distribution. The Multiple Random Eigengraphs (MREG) is a mathematical framework modeling multiple random graphs(Wang et al., 2019). Consider a set of *m* unweighted and undirected graphs with the same *n* vertices denoted by 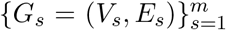. Let *h*_1_,…,*h_d_* be normalized vectors in 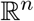 that span a subspace of dimension *d*, contributing to a large amount of variability in the set of graphs, and *λ*_1_,…, *λ_d_* be vectors in 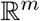 such that 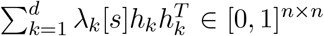 for all *λ*(Wang et al., 2019) for subject *s*. Then the adjacency matrix *A_s_* for each graph *G_s_* should be modeled as follows(Wang et al., 2019):

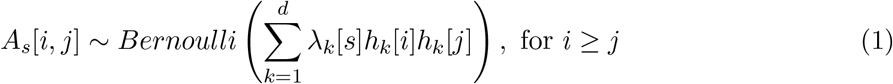

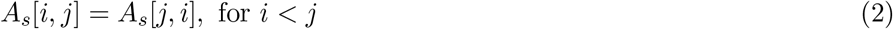

Note here each *A_s_* is symmetric, as opposed to sampling independently above and below the diagonal. The *h* vectors span the latent subspace shared by the set of multiple graphs, and the *λ* vectors represent graph-specific loadings onto the subspace(Wang et al., 2019).

The original joint graph embedding algorithm by Wang et al.(Wang et al., 2019) estimates a low dimensional embedding of the latent space across multiple graphs and each graph’s projection onto that embedding under the MREG model. It estimates the subspace by minimizing the sum of squared errors (SSE) between the subspace and adjacency matrices.(Wang et al., 2019). In this work, we implemented a modified version of the Wang et al. algorithm as discussed below.

### 2.3 Limitations of the original framework

There are several limitations to consider when applying the original framework and algorithm above to neurodegeneration data, which we will address and modify in our version:

1. The constraint that probabilities lie in [0,1] is difficult to enforce in practice
2. Incorrect estimation of diagonal entries contributes a negligible amount of error when the number of vertices is large, but contributes significantly in our case.
3. The SSE loss function does not correspond to a log likelihood under the proposed Bernoulli model, and therefore resulting parameter estimates do not have desirable properties of maximum likelihood estimators

Now we state our main contribution and novelty in this work. First, we address the three limitations listed above by (1) introducing a sigmoid function to the model to guarantee edge probabilities are in [0, 1], (2) adding constraints on the diagonal such that parameter estimates are not forced to fit diagonal entries that carry no meaning (since diagonal entries of a correlation matrix are always 1), and (3) extending the original algorithm(Wang et al., 2019) to identify maximum likelihood estimators by gradient descent. With these modifications, our embedding algorithm generates probabilities suitable for a Bernoulli model, which we will describe more in detail below. Secondly, we develop and implement a novel statistical testing framework to detect complex patterns including network-structure pairs and triples, rather than a machine learning classifier. To our knowledge, our approach to testing patterns has not been performed to analyze joint graph embedding results on brain imaging data before.

### 2.4 Modifications to MREG

In this section, we precisely state our modifications to the MREG model in Wang et al.(Wang et al., 2019). Consider the set of *m* unweighted and undirected graphs with the same *n* vertices denoted by 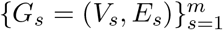, where a vertex represents a brain structure of our interest and an edge represents a strong correlation between structures. We modify the interpretation of *h* and state that the *h* vectors now span a space of parameters that encode the probability when acted on by a sigmoid function. This sigmoid function guarantees 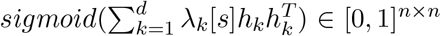 even in the case of small samples, addressing limitation 1. Secondly, we force diagonal entries to be 1 since a structure’s relation with itself is not of interest in this work, addressing limitation 2. Our modifications to equation 1 and 2 are as follows:

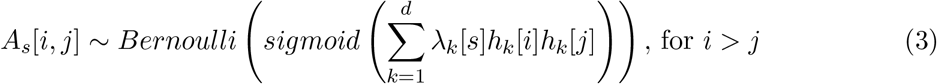

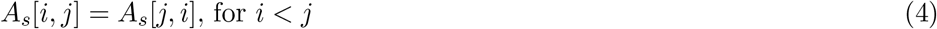

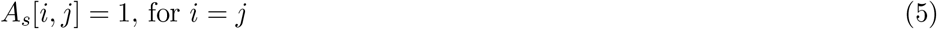

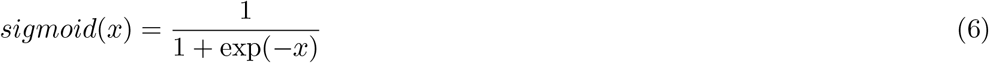

An example transformation from the latent subspace and subject specific loading to adjacency matrices observed in our data is illustrated in Fig. 2.

**Fig. 2.**
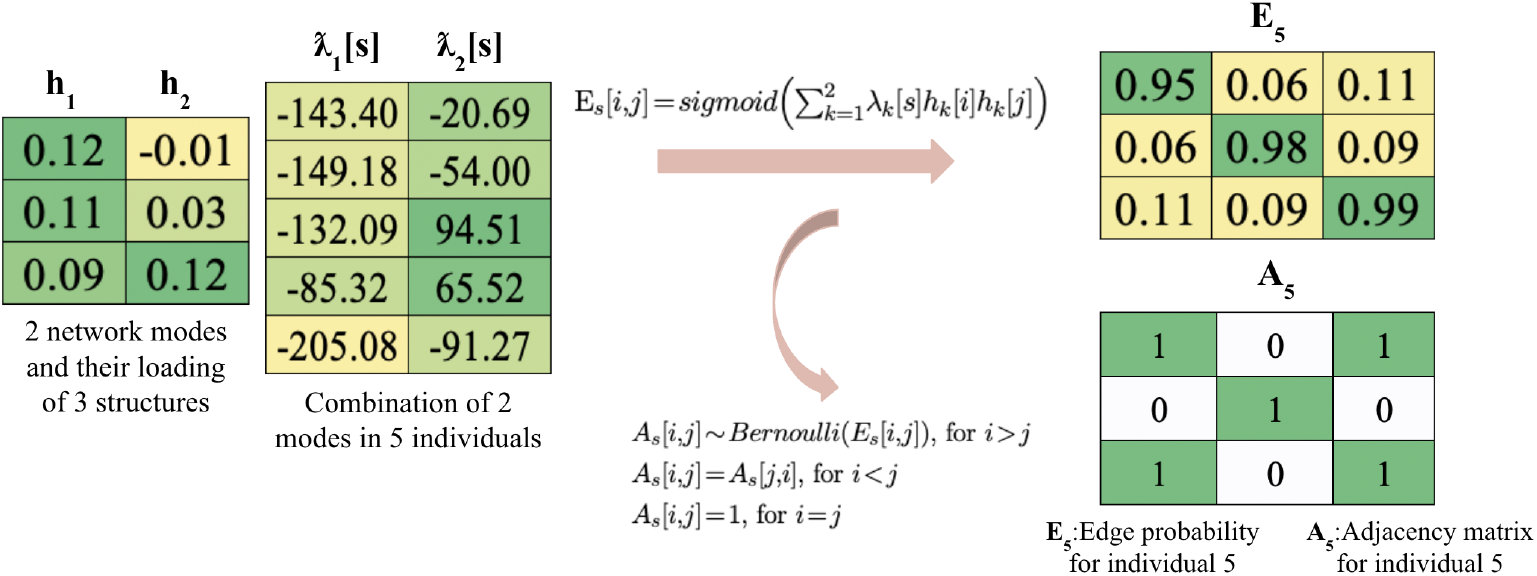
An example of MREG for the same individual and structure in Fig. 1

### 2.5 Our Maximum Likelihood Joint Graph Embedding Algorithm

We change the algorithm to better fit the constraints of our study and estimate the subspace by minimizing the binary cross entropy (BCE) loss, which is the log likelihood under the model. Let *E_s_* [*i,j*] be a symmetric matrix representing edge probabilities between structures *i* and *j* for subject *s*, then the BCE loss is minimized as in equation (7), where *A_s_* are the observed adjacency matrices:

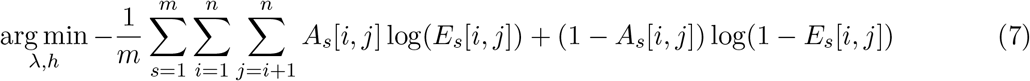

Note here that the diagonal terms do not contribute to our loss function. By replacing the original loss function with BCE, we will have a well characterized maximum likelihood estimator even in the small sample case, addressing limitation 3. We take a greedy approach in finding the optimal representation of the latent space, where we start with a 1D optimization problem, and then expand to the second dimension while keeping the first dimension fixed to find the optimal representation. The algorithm is implemented as follows:

~~~
  *λ* ← random.normal((m, number of networks)), *h* ← random.normal((n, number of networks)),
  normalize *h*
  **for** *d* = 1; *d* <=number of networks; *d* + + **do**
      **while** loss not coverged in dimension *d* **do**
            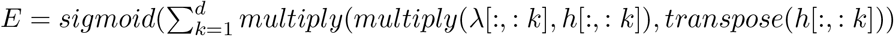
            Compute BCE loss as in equation (7)
            Compute gradient of the BCE with respect to *λ* and *h*
           Update *λ*[:,*d*] by gradient descent, and *h*[:,*d*] by projected gradient descent (i.e. update
  and
            normalize)
     **end while
  end for**
~~~

We coded in python and PyTorch, and used automatically calculated gradients for gradient descent. The algorithm took 30 minutes to run on a computer with 10-core CPU. Convergence in all 4 dimensions is shown in Fig. 3a, where at every 10,000 iterations we observe the loss dropping quickly after adding another dimension, and converging before the next dimension is added.

**Fig. 3.**
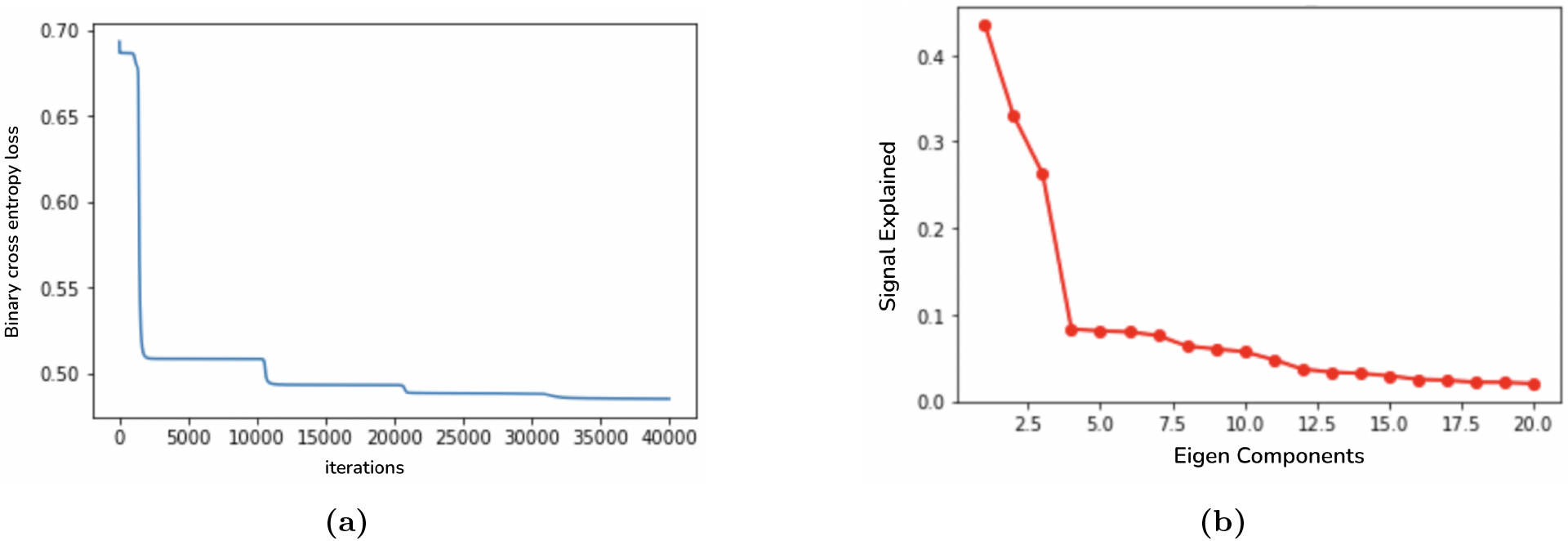
a). Convergence in all 4 dimensions with the algorithm b). Scree plot with elbow at 4 dimensions

### 2.6 Hyperparameter Tuning

For hyperparameter tuning, we first estimate an optimal dimension of the latent space to account for the majority of the variation in observed correlation matrices. To do this, we computed an eigendecomposition for each subject’s correlation matrix, and found that 4 components provided a reasonable reconstruction accuracy. A scree plot for one typical subject is shown in Fig. 3b. Next, we performed a grid search over several orders of magnitude for the gradient descent step sizes corresponding to *h* and *λ*, and selected the largest parameters that gave convergence without oscillation. We decided to set the step size for *h* to 2 and the step size for *λ* to 20000.

### 2.7 Significance Testing

We develop and implement a novel test statistic similar to F-type statistics (i.e. comparing sum of square error under two different models) to test for differences in networks and structures between the two groups, and use permutation testing on a maximum statistic to control for Familywise Error Rate (FWER) at 5% (Nichols and Hayasaka, 2003). In addition, we compare the difference between groups after accounting for confounders: age, intracranial volume and APOE gene status by least squares regression. APOE gene status was modeled as a categorical (as opposed to cardinal) variable, where an individual may have 0, 1, or 2 copies. For a cohort with *m* subjects, *d* networks, and *n* structures, we define the test score as 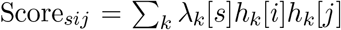 for subject s and structures *i* and *j*. We pass the score through a sigmoid to give probabilities *E*, and estimate these variables by maximum likelihood. We form the tensor *T_skij_* = *λ_k_*[*s*]*h_k_*[*i*]*h_k_*[*j*] to get a three-tuple for each subject *s*, of which we will perform statistical tests by comparing how close it is to its group-dependent or group-combined average, after accounting for confounders. By summing over various combinations of indices in *T* before statistical testing, we are able to test for:

1. networks only
2. structures only
3. network-structure pairs
4. structure-structure pairs
5. network-structure-structure triples

As other standard tests for items 2 and 4 exist, we will focus our work on 1, 3, and 5. To our knowledge, our framework for testing patterns involving networks, pairs, or triples has not been performed to analyze brain imaging data before.

#### 2.7.1 Confounder regression analysis

We perform least square regression analysis to test for true signals not caused by common confounders for AD and neurodegeneration. We start with a design matrix *D* containing a column of 1s (for mean), and columns for age, intracranial volume and APOE gene status, and estimate a coefficient matrix 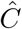 for confounders. From 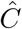 the SSE for one group is calculated as follows:

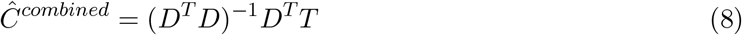

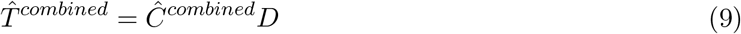

Here *T* is reshaped into a matrix from a tensor for calculation. Next, we calculate the SSE for splitting the cohort into two groups: none/mild and severe AD. We first form *D’*, which has an additional column to *D* indicating disease status. We then perform the same regression, with *D’* replacing *D* in equations 8 and 9 to find 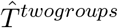. In the next sections, we will use 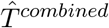 and 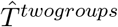 to calculate the sum of square errors (SSE) when considering one vs. two groups.

#### 2.7.2 Networks

We start by testing for significant networks and obtain the test statistic *X_k_* for network *k* by reducing (i.e. summing over) additional dimensions. Let 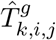 be the expected value matrix under our linear model for dimension *k*, structures *i* and *j*, and subjects in group *g*. We remove extra dimensions by calculating the SSE between one group vs. two groups and *X_k_* as follows:

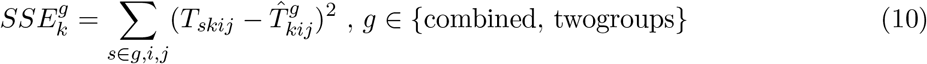

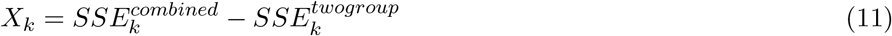

To control the FWER at 5%, we use permutation testing and take the max over *k* at each iteration and define the threshold as the 95 percentile of 10,000 simulation results(Nichols and Hayasaka, 2003).

#### 2.7.3 Network structure pairs

Similar to networks only, we calculate the SSE in the two settings but reduce one fewer level as follows:

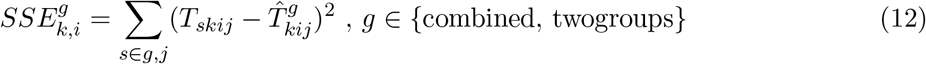

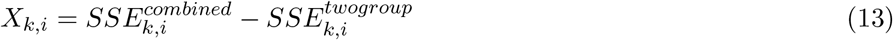

We follow the same permutation testing procedure and take a max over *k*, *i* instead of just *k* at each iteration.

#### 2.7.4 Network structure structure triples

Lastly for triples, we form the test statistic:

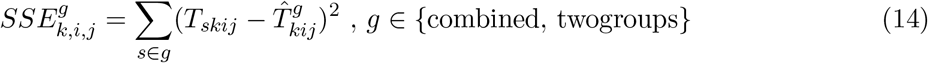

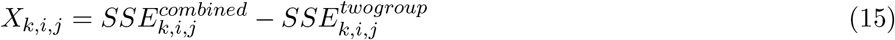

We follow the same permutation testing procedure and take a max over *k, i, j* at each iteration.

In our experiments, we performed the analysis twice: with and without adjusting for confounders. Note that not adjusting for confounders is a special case of *T* estimation, where 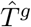 is simply the mean over all subjects in group *g*. In the results section, we will present the former, and make brief comparisons to the latter.

### 2.8 Code availability

Our source code and documentation are available on GitHub at https://github.com/twardlab/joint_graph_embedding_AD. To replicate our work, first agree to the user agreement by ADNI and download the ADNI1 3Y1.5T Longitudinal FreeSurfer dataset by University of California, San Francisco(Wyman et al., 2013). Put all files under a directory named dataset, and first run the Jupyter notebook *preprocess*, then the notebook *joint-graphembedding-analysis*. We document our code with Sphinx(Brandl, 2021), and save documentations in *docs*. More details on how to replicate our work can be found in our GitHub repository.

## 3 Results

### 3.1 Significant networks associated with AD neurodegeneration

Out of the 4 networks identified from joint graph embedding, we found the first 2 extremely significant, after accounting for confounders. Upon examination in Table 1, both networks are dominated by structures believed to be associated with AD neurodegeneration. We show the results from permutation testing and the statistics for each network in Fig. 4 and visualize the structures in network 1 and 2 in Fig. 5. In Table 1, we include the top 10 structure by absolute value of loading in each network.

**Fig. 4.**
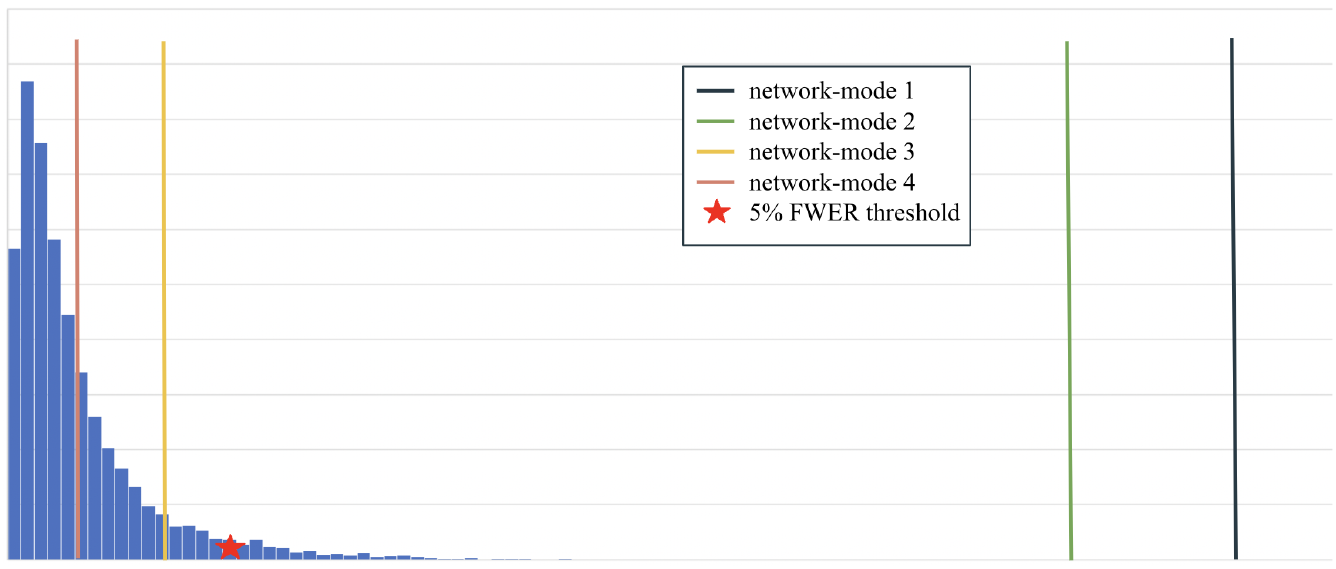
Histogram of permutation testing after accounting for confounders

**Fig. 5.**
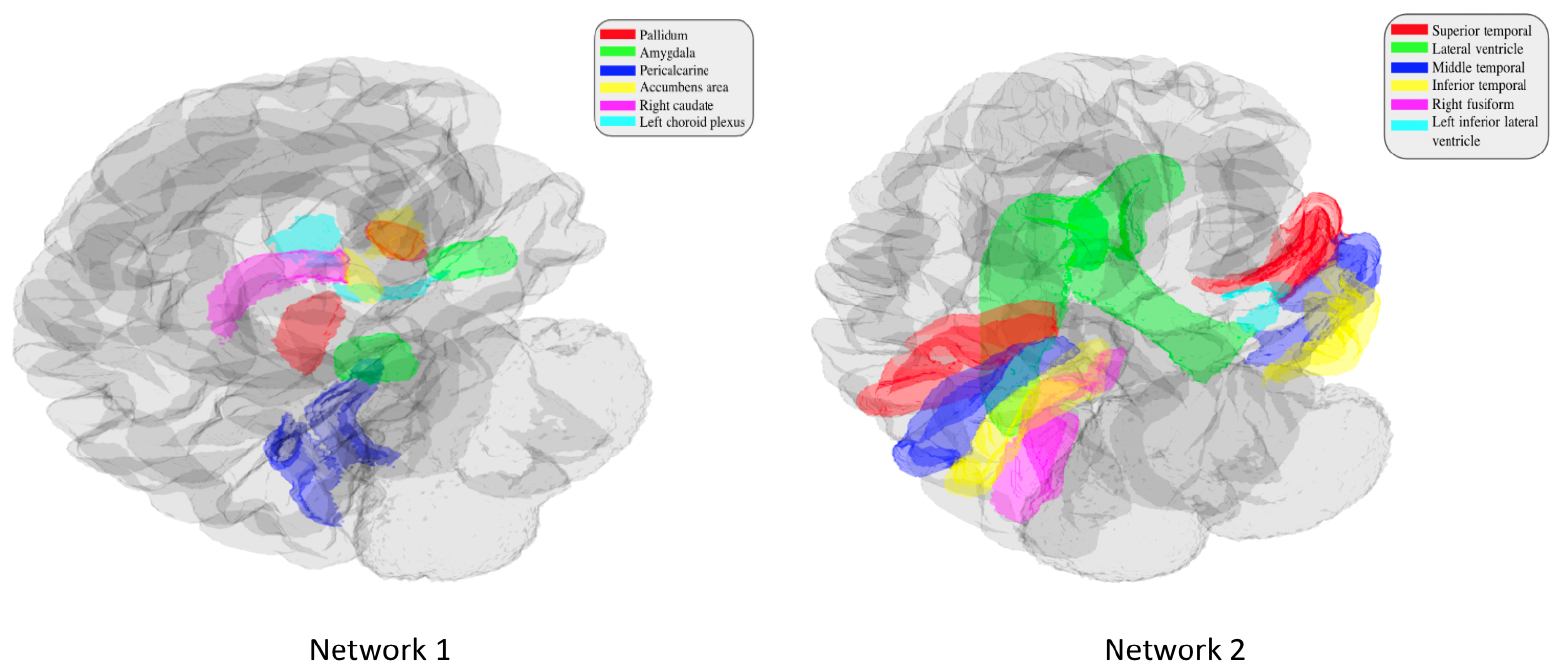
Visualization of significant networks

**Table 1.** Top 10 structures ranked by loading of each network

| network 1 | network 2 | network 3 | network 4 |
| --- | --- | --- | --- |
| Left Pericalcarine | Right Middle Temporal | Right Caudal Middle Frontal | Fifth Ventricle |
| Left Pallidum | Left Middle Temporal | Right Superior Frontal | Right Lingual |
| Right Pericalcarine | Left Inferior Temporal | Right Rostral Middle Frontal | Left Precuneus |
| Right Pallidum | Left Superior Temporal | Left Superior Frontal | Right Lateral Occipital |
| Right Accumbens Area | Right Inferior Temporal | Left Caudal Middle Frontal | Left Superior Parietal |
| Right Amygdala | Right Lateral Ventricle | Left Rostral Middle Frontal | Right Precuneus |
| Right Caudate | Left Lateral Ventricle | Right Inferior Parietal | Right Superior Parietal |
| Left Accumbens Area | Right Superior Temporal | Left Precentral | Left Inferior Parietal |
| Left Choroid Plexus | Left Inferior Lateral Ventricle | Right Precentral | Left Isthmus Cingulate |
| Left Amygdala | Right Fusiform | Right Supramarginal | Left Cerebral Cortex |

### 3.2 Significant network structure pairs associated with AD neurodegeneration

For network-structure pairs, we rejected 170 out of 432 network structure pairs. While most rejected pairs occur in networks previously found significant, the pairs are not exclusive to network 1 and 2 only. Three structure pairs in network 3 were found significant. In Table 2, we show the top 5 significant pairs ranked by p-value.

**Table 2.** Top 5 significant pairs ranked by FWER-corrected p value

| network-structure pairs | p-value |
| --- | --- |
| 1-Right Pallidum | <1e-4 |
| 2-Left Inferior Lateral Ventricle | <1e-4 |
| 2-Left Fusiform | <1e-4 |
| 2-Left Entorhinal | <1e-4 |
| 2-Left Cerebral Cortex | 1e-4 |

### 3.3 Significant network structures triples associated with AD neurodegeneration

For network-structure-structure triples, we rejected 753 out of 46656 possible triples. The network and structures found significant are again not a subset of those found significant in the network-structure pair. In Table 3, we show the top 5 significant triples ranked by p-value.

**Table 3.** Top 5 significant triples ranked by FWER-corrected p value

| network-structure-structure triples | p-value |
| --- | --- |
| 2-Left Inferior Temporal-Right Middle Temporal | <1e-4 |
| 2-Left Middle Temporal-Right Middle Temporal | <1e-4 |
| 2-Left Superior Temporal-Right Middle Temporal | <1e-4 |
| 2-Left Inferior Temporal-Left Middle Temporal | <1e-4 |
| 2-Left Middle Temporal-Left Superior Temporal | 1e-4 |

### 3.4 Results comparing confounder regression and no regression

While the results were similar to those shown above, the analysis without accounting for confounders found more structures and networks significant than analysis with confounder regression. For networks only, network 3 became slightly significant, and networks 1 and 2 remained highly significant. The top 10 structures were the same for networks 1 and 2 in both analysis, but those in structure 3 and 4 were different. For tuples analysis, we found 205 instead of 170 pairs and 1171 instead of 753 triples significant. The networks and structures found significant in triples are not a subset of those found in pairs and vice versa.

## 4 Discussion

In this work, we applied a joint graph embedding method (similar to a multivoxel dictionary embedding method), which would typically be used to study functional connectivity, to volumetric data in neurodegeneration. We extended the original algorithm(Wang et al., 2019) and implemented our maximum likelihood joint graph embedding algorithm to identify significant structural networks from volumetric data in Alzheimer’s disease cohorts(Wyman et al., 2013). We showed that our version of the algorithm has promise in uncovering latent dimensions that are easy to interpret and visualize. In addition, we developed and implemented a novel testing procedure and tested for significant networks, network-structure pairs and network-structure-structure triples. Since these pairs and triples are a novel description of neurodegeneration patterns, we will briefly state their interpretation. For example, our discovered pairs can be interpreted as “the right pallidum displays a significant association with disease status, through its role in network 1”. As another example, our discovered triples can be interpreted as “the interaction between the left inferior temporal lobe and right middle temporal lobe displays a significant association with disease status, through its role in network 2”. We performed analysis to regress out common confounders in AD in hope of gaining more discovery power and make a few comments here between our results. We found fewer significant pairs and triples when taking confounders into account than not. This is expected, as some structures previously found significant may be caused by confounders. While networks 1 and 2 remained unchanged and highly significant, indicating that changes in these networks are due to disease progression, network 3 was no longer a significant network. The structures’ change in networks 3 and 4 also indicate these structures may have been significant due to common confounders such as age.

Next we discuss a few limitations in this work. First, due to our high standard of cohort selection, this dataset may not be representative of clinical MRI subjects. As such, a natural future direction for the study is to apply this framework to clinical datasets, which represent a more diverse population. Secondly, our findings are on a population scale instead of an individual scale, which may lead to additional biases when applying to individual clinical diagnosis. We believe that using a more fine-grained separation of the cohort, as opposed to modeling disease status as only two groups, may help address this issue. Thirdly, we note that since there has been little work done to apply graph embedding methods to study volumetric data in Alzheimer’s Disease cohorts, we do not have result comparisons with “state-of-the-art” methods. As with most unsupervised methods, there is no ground truth for evaluation to draw a conclusion on which method is best. Rather, we offer an additional method in analyzing group differences between healthy and diseased individuals based on neurodegeneration.

Our framework shows promise in that it discovered structures commonly believed to be associated with AD neurodegeneration. We point out several strengths in our framework. First, each of the three groups of findings gives us more information on structural correlations than traditional methods(Pengas et al., 2012), and we also find structures not found by previous methods, such as mass univariate analysis(Bernal-Rusiel et al., 2013). Secondly, the dataset used in our study, ADNI(Jack Jr et al., 2008), is very well characterized, it is more robust and spans a longer period than most clinical data. Thirdly, though increased complexity in algorithm and testing procedure often require more computational time, our algorithm embeds 494 patients with 108 structures in just 30 minutes and runs significance testing in 4 hours. We experimented with cohort and structure sizes and show runtime results in Fig. 6. We note that while the algorithm’s runtime scales linearly in subjects, it is exponential in structure counts. However, as studies often do not include a large set of structures, it should not be of big concern to users. On the other hand, our algorithm shows good time complexity with increased cohort sizes. By offering more flexibility in terms of patterns that we can identify, we hope that our framework will reveal biomarker patterns more sensitive to AD.

**Fig. 6.**
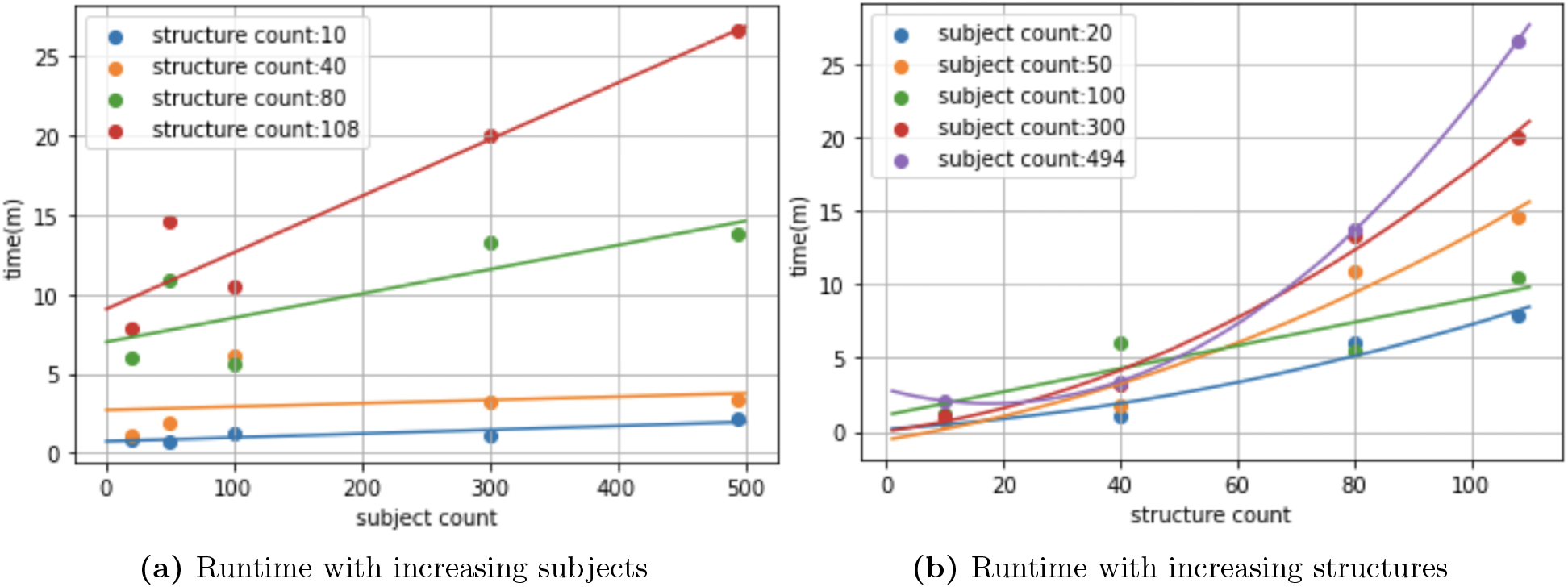
Runtime with increasing parameters

With the recent development of potential drugs to treat Alzheimer’s disease, methods like ours that can quantify complex patterns of neurodegeneration will be essential for noninvasively identifying patients who would benefit the most. We believe that the development and dissemination of algorithms such as this one, through open source code and well documented examples, will play an important role in helping to reduce the burden of Alzheimer’s disease on our aging population.

## Information Sharing Statement

Our code is publicly available on Github, details on access in section 2.8. The sample data can be downloaded from ADNI after accepting their data use agreement.

## Acknowledgement

Data collection and sharing for this project was funded by the Alzheimer’s Disease Neuroimaging Initiative (ADNI) (National Institutes of Health Grant U01 AG024904) and DOD ADNI (Department of Defense award number W81XWH-12-2-0012). ADNI is funded by the National Institute on Aging, the National Institute of Biomedical Imaging and Bioengineering, and through generous contributions from the following: AbbVie, Alzheimer’s Association; Alzheimer’s Drug Discovery Foundation; Araclon Biotech; BioClinica, Inc.; Biogen; Bristol-Myers Squibb Company; CereSpir, Inc.; Cogstate; Eisai Inc.; Elan Pharmaceuticals, Inc.; Eli Lilly and Company; EuroImmun; F. Hoffmann-La Roche Ltd and its affiliated company Genentech, Inc.; Fujirebio; GE Healthcare; IXICO Ltd.;Janssen Alzheimer Immunotherapy Research & Development, LLC.; Johnson & Johnson Pharmaceutical Research & Development LLC.; Lumosity; Lundbeck; Merck & Co., Inc.;Meso Scale Diagnostics, LLC.; NeuroRx Research; Neurotrack Technologies; Novartis Pharmaceuticals Corporation; Pfizer Inc.; Piramal Imaging; Servier; Takeda Pharmaceutical Company; and Transition Therapeutics. The Canadian Institutes of Health Research is providing funds to support ADNI clinical sites in Canada. Private sector contributions are facilitated by the Foundation for the National Institutes of Health (www.fnih.org). The grantee organization is the Northern California Institute for Research and Education, and the study is coordinated by the Alzheimer’s Therapeutic Research Institute at the University of Southern California. ADNI data are disseminated by the Laboratory for Neuro Imaging at the University of Southern California.

This work was supported by The Karen Toffler Charitable Trust through the Toffler Scholars Program. Funders played no role in study design; in the collection, analysis and interpretation of data; in the writing of the report; or in the decision to submit the article for publication.

